# Piezo1 activation attenuates thrombin-induced blebbing in breast cancer cells

**DOI:** 10.1101/2020.04.30.068338

**Authors:** Paul O’Callaghan, Adam Engberg, Nikos Fatsis-Kavalopoulos, Gonzalo Sanchez, Olof Idevall-Hagren, Johan Kreuger

## Abstract

Cancer cells exploit a variety of migration modes to leave primary tumors and establish metastases, including amoeboid cell migration facilitated by bleb formation. Here we demonstrate that thrombin induces dynamic blebbing in the MDA-MB-231 breast cancer cell line, and confirm that PAR2 activation is sufficient to induce this effect. Cell confinement has been implicated as a driving force in bleb-based migration. Unexpectedly, we find that gentle contact compression, exerted using a “Cell Press” to mechanically stimulate cells, attenuated thrombin-induced blebbing with an associated increase in cytosolic calcium. Thrombin-induced blebbing was similarly attenuated using Yoda1, an agonist of the mechanosensitive calcium channel Piezo1, and its capacity to attenuate blebbing was impaired in Piezo1 depleted cells. Additionally, Piezo1 activation suppressed and reversed the thrombin-induced phosphorylation of ERM proteins, which are implicated in the blebbing process, and this activity was in part mediated through activation of the PP1A/PP2A family of serine/threonine phosphatases. Our results provide mechanistic insights into Piezo1 activation as a suppressor of dynamic blebbing, specifically that which is induced by thrombin.

## Introduction

Cancer cells adopt a variety of migratory modes during metastasis, influenced in part by the nature of the extracellular matrix surrounding them^1,2^. The amoeboid mode of migration is defined by the pressure-driven expansion and actomyosin-mediated retraction of plasma membrane (PM) protrusions, termed blebs. Bleb-driven cell migration is observed in a number of *in vivo* settings including germ cell migration during zebrafish embryogenesis^3,4^ and in human cancer cells in models of tumorigenesis^1^. Bleb formation involves an expansion of the PM, triggered by its dissociation from the underlying actin cortex, causing it to balloon outward due to hydrostatic pressure^5^. Ezrin, radixin and moesin (ERM) proteins, which tether actin to the PM, are among the first components recruited to the cytoskeletal-free bleb membrane, followed by actin and then myosin, which completes assembly of the contractile machinery required to retract the bleb membrane^6,7^. The potential for mechanical confinement to trigger cells to adopt blebbing phenotypes and amoeboid modes of migration has been extensively studied and provides a basis for cell movement through matrices with reduced dependence on focal adhesions^1,8–10^.

Tumor vascularization is central to the expansion of the primary malignancy, but is also the conduit through which cancer cells escape and establish metastases. Tumor vessels are notoriously leaky and the tumor environment has been described as being in a pathological state of coagulation, which may be exacerbated by certain chemotherapies^11^. Consequently, cancer cells may exist within, and contribute to, a milieu enriched in various components of the coagulation cascade, including thrombin^12^ (reviewed in ^13^). Thrombin acts on the G protein-coupled protease activated receptors (PARs) 1-4^14–18^ to irreversibly sever a part of their N-terminal extracellular domain exposing a tethered ligand, which activates the receptor through an intramolecular interaction. Thrombin activation of PARs results in phospholipase-C β (PLCβ)-mediated release of inositol 1,4,5-trisphosphate (IP3) from phosphatidylinositol 4,5-bisphosphate (PIP2), leaving diacylglycerol (DAG) in the PM, which can bind protein kinase C (PKC). IP3 binding to IP3 receptors (IP3R) in the endoplasmic reticulum (ER) triggers release of stored Ca^2+^, which in turn activates the DAG-bound PKC (reviewed in^19^). Ca^2+^ acts in consort with a variety of binding partners and relevant here is the Ca^2+^/calmodulin (CaM)-mediated activation of myosin light chain kinase (MLCK), which phosphorylates myosin II permitting it to bind and contract actin filaments (reviewed in^20^). Myosin activity is fundamental to the contractility of cellular cytoskeletons and actomyosin contraction is an established driver of blebbing through its capacity to increase hydrostatic pressure within the cell^21^; in parallel, actomyosin contractility is essential for the retraction of expanded PM blebs^5^.

Piezo1 is a mechanosensitive cation channel^22,23^ implicated in diverse areas of cell biology including shear stress sensing in endothelial cells^24^; regulation of red blood cell volume^25^; cell division in epithelial cells^26^; and as a confinement sensor that suppresses PKA activity to optimize cell motility^27^. Piezo1 is found in the plasma membrane as a homotrimer and in closed conformation induces a bowl-shaped indentation in the PM. In response to increased membrane tension Piezo1 transitions to a more planar arrangement and gates Ca^2+^ influx^28,29^. Furthermore, loss of cortical actin, as seen during the initial expansion of PM blebs, reduces the activation threshold of Piezo1^30^.

Here, we identify thrombin as a bleb induction stimulus, and Piezo1 activation, through contact compression or stimulation with the Piezo1 agonist Yoda1, as an effective mechanism to suppress dynamic blebbing in breast cancer cells. Thrombin induced blebbing was associated with increased ERM phosphorylation that was suppressed or reversed through Piezo1 activation, which in turn was dependent on the downstream activity of PP1/PP2A serine/threonine phosphatases. We propose that Ca^2+^ influx via Piezo1 activates PP1/PP2A phosphatases to dephosphorylate ERMs, which in turn reduces the actomyosin contraction forces applied to the PM; thereby reducing the likelihood of dynamic blebbing.

## Materials and Methods

### Reagents and chemicals

SiR-Actin (Spirochrome, Tebu-bio, Roskilde, Denmark) was diluted to 1 mM in dimethyl sulfoxide (DMSO; ThermoFisher Scientific, Uppsala, Sweden) and stored at −20 °C. Fluo-4 and Fura Red calcium indicators (ThermoFisher Scientific) were resuspended in DMSO to 1 mM prior to dilution and cell staining. Human alpha thrombin (Nordic Diagnostica AB, Billdal, Sweden) was resuspended in OptiMEM reduced serum medium (RSM) (Thermo Fisher Scientific) to 1 U/μl and stored at −20 °C. PAR2 agonist peptide SLIGRL-NH2 (Abcam, Cambridge, UK) was diluted to 1 mM in OptiMEM RSM and stored at −20 ° C. Yoda1(Tocris, Bio-Techne, Abingdon, UK), the Piezo1 channel activator, was diluted to 20 mM in DMSO and stored at −20°C. GsMTx-4, the mechanosensitive and stretch-activated ion channel inhibitor (Abcam, Cambridge, United Kingdom), was diluted to 250 μM in DMSO and stored at −20°C. Calyculin A (Tocris) was diluted to 1 mM and Cyclosporin A (Tocris) was diluted to 100 mM in DMSO and stored at −20°C. Polydimethylsiloxane (PDMS) pillars (Sylgard 184 silicone elastomer using a 10 (base):1 (curing agent) ratio (Sigma-Aldrich Sweden AB, Stockholm, Sweden)) were cast from a custom mold produced with a Form2 3D printer (Formlabs).

### Antibodies

Piezo1 NBP1-78446 rabbit pAb (Novus Biologicals, Bio-Techne, Abingdon, UK) was detected with Alexa Fluor Plus goat anti-rabbit IgG secondary antibodies conjugated to 488 nm fluorophores, for Piezo1 immunocytochemistry. Phospho-ezrin (Thr567)/radixin (Thr564)/moesin (Thr558) mouse mAb 48G2 (Cell Signaling Technology) and alpha (α)-tubulin (11H19) rabbit mAb (Cell Signaling Technology) were detected with Alexa Fluor goat anti-rabbit IgG secondary antibodies conjugated to 680 nm fluorophores nm, for phospho-ERM and a-tubulin immunoblotting.

### Cell culture

The MDA-MB-231 breast cancer cell line was obtained from ATCC (LGC Standards GmbH, Wesel, Germany) and externally authenticated using Idexx Bioanalytics (Ludwigsburg, Germany), which confirmed their genetic signature to be consistent with that published for the cell line of origin^31^. Cells were cultured in DMEM Glutamax supplemented with 10% FBS (Thermo Fisher Scientific, Uppsala, Sweden) under standard conditions of 37°C, 5% CO_2_.

### Spontaneous blebbing assessment

MDA-MB-231 cell cultures were maintained in DMEM Glutamax, 10% FBS and imaged using a 20x objective and a combination of confocal and DIC microscopy. Time-lapse sequences of cells were analyzed in ImageJ, and individual cells in each field of view were manually annotated as regions of interest (ROI) using the polygon selection tool to facilitate total cell counting. Each ROI was visually assessed for the duration of the time-lapse sequence and blebbing cells were manually annotated with the ImageJ point tool. The percentage of blebbing cells per field of view was calculated in Microsoft Excel.

### Live cell imaging and treatments

For all live-imaging experiments cells were seeded in MatTek (MatTek Corporation, Bratislava, Slovak Republic) or Ibidi (LRI Instrument Ab, Lund, Sweden) 35 mm dishes with uncoated coverslip bottoms. Prior to live imaging of F-actin, cells were incubated overnight with 0.5 μM SiR-actin in DMEM Glutamax, 10% FBS. For calcium imaging, cells were incubated with 1 μM Fluo-4 and Fura Red in OptiMEM RSM, without phenol red (referred to hereafter as OptiMEM) for 1 h, after which cells were washed and cultured for an additional 1.5 h in OptiMEM before imaging and treatments. Time-lapse imaging of Fluo-4 and Fura Red fluorescence, and differential interference contrast (DIC) channels was performed on an LSM700 confocal microscope (Zeiss, Jena, Germany) equipped with an incubation chamber and heated stage, using a Plan-Apochromat 20x/0.8 (Zeiss) objective with a pinhole setting of 1 Airy unit. Fluo-4 fluorescence and DIC images were scanned simultaneously on track 1, followed by Fura Red fluorescence scanning on track 2. The total acquisition time for one frame consisting of Fluo-4, Fura Red and DIC signals in the time-lapse imaging experiments was approximately 9.2 s. Prior to the indicated treatments a baseline for Fluo-4 and Fura Red fluorescence was established. Treatments were administered manually by pipetting reagents directly into the culture dish. To facilitate access to the culture dish when administering treatment solutions, the head of the microscope was tilted backwards, which temporarily obstructed DIC imaging. The stated treatment concentrations account for the dilution of reagents in the imaging medium.

### Image analysis and processing

Time-lapse sequences of Fluo-4 and Fura Red fluorescence and DIC recordings were analyzed using ImageJ. Individual cells in the DIC recordings were viewed for the duration of the time-lapse experiment and the total area occupied by each cell was outlined with the polygon tool and annotated as a region of interest (ROI). The Fluo-4 and Fura Red fluorescence intensity was measured for each ROI using the multi-measure function in ImageJ and the values were exported to Excel where the Fluo-4/Fura Red ratio, the relative Fluo-4/Fura Red signals, and the maximum and accumulated Fluo-4/Fura Red responses were calculated for the durations of treatments indicated in the respective figures. The blebbing status for each cell during the indicated treatments was visually assessed and manually recorded in Excel as not blebbing (NB); blebbing with small less well defined blebs (sB); or blebbing with well-defined blebs (B). Where it was difficult to visually discern the boundaries for entangled clusters of 3-4 cells the area occupied by the cluster was annotated as a single ROI for the purpose of Fluo-4 and Fura Red fluorescence analysis, but where possible the blebbing status for each cell in the cluster was individually recorded. In the comparison of the Fluo-4/Fura Red signal in siPiezo1- and siCtrl-treated cells, the maximum response to thrombin was arbitrarily set to 100% for each experiment to permit a relative comparison of the Yoda1 response between siPiezo1 and siCtrl groups. Kymographs of DIC and Fluo-4/Fura Red recordings were prepared by applying the ImageJ reslice command to linear ROIs plotted through individual cells. To prepare the ratiometric Fluo-4/Fura Red kymographs using the ImageJ Image Calculator tool the boundary defined by the Fluo-4 and Fura Red signals in a composite image of the resliced cell was assigned as an ROI. The ImageJ Clear Outside function was used to clear fluorescence from all pixels outside of this ROI (i.e. in the cell free background). The Fluo-4 signal was then divided by the Fura Red signal and the ratiometric fluorescence was presented; notably, no quantitative analysis was performed on images processed like this.

### Cell compression

Cell compression experiments were conducted with the “Cell Press”: a custom-built manipulator composed of 3D printed parts; a piezoelectric positioner (SmarAct GmbH, Oldenburg, Germany) operated with an MCS2 manual controller (SmarAct); and a flexible PDMS pillar cast from a 3D printed mold. The Cell Press was fixed to the microscope behind the translation stage (Fig. 3a). To control the position of the PDMS pillar relative to the cell layer we established and recorded the z-position displayed on the MCS2 controller for the point-of-contact between the surface of the PDMS pillar and the bottom of a cell-free dish as follows: Ink markings were made on the surface of the pillar and the surface of the coverslip. The piezoelectric positioner was moved to its start position ensuring the pillar was at its maximum distance from the coverslip. While visualizing the ink marking on the coverslip under the microscope the pillar was lowered until its ink marking was also visible. The positioner was then set to descend in 5 μm steps. We observed that, due to its flexibility, once the PDMS pillar was in contact with the coverslip it moved in synch with small x, y positional changes to the coverslip. Therefore, for each step the pillar was lowered we tested if it had made contact with the coverslip by making small adjustments to the translation table until both ink markings moved in synch, indicating that the pillar was in contact with the coverslip. This position was recorded and used as a reference point for subsequent cell compression experiments. Fluo-4- and Fura Red-labelled cells were visualized as before and the pillar was then lowered until it was within approximately 100 μm of the recorded reference point, permitting a baseline of non-compressed signal to be recorded. The positioner was then lowered in 5 μm steps until a calcium response was observed, this was considered the point of contact compression (CC). For deformation compression (DC) the pillar was further lowered until the cells were visibly deformed.

### Piezo1 knock down

MDA-MB-231 cells were co-transfected with Silencer Select pre-designed siRNAs si18891 and si18892, which respectively target the sequences GCCTCGTGGTCTACAAGATT and AGAAGAAGATCGTCAAGTA of human Piezo1 mRNA. siRNAs (siPiezo1 and siCtrl) were delivered using Lipofectamine RNAiMax transfection solution (ThermoFisher Scientific). Experiments were performed 48 h post-transfection.

### Reverse transcription and real-time qPCR

RNA was isolated from siPiezo1- or siCtrl-transfected MDA-MB-231 cells using a PureLink RNA Mini Kit (Thermo Fisher Scientific), concentrations were determined by NanoDrop (Thermo Fisher Scientific). cDNA was synthesized by reverse transcription using the iScript kit (BioRad, Stockholm, Sweden). Transcripts of interest were analyzed using the QuantStudio 5 real-time quantitative PCR (RT-qPCR) system with the associated software (Thermo Fisher Scientific). cDNA was diluted with the Fast SYBR green master mix (Thermo Fisher Scientific) and Piezo1 mRNA levels were assessed by RT-qPCR using the following primer pairs: forward CACCAACCTCATCAGCGACT and reverse GCACCAGCCAGAACAGGTAT; and forward AGGGAGGCACTGTGGAGTAT and reverse AGGGATGACCACAGACTGGT. Similarly, Piezo2 mRNA levels were assessed using the following primer pairs: forward CACCATCTACAGACTGGCCCAC and reverse ACCAGGTGCCATTTGTTCCTT; and forward GACGGACACAACTTTGAGCCTG and reverse CTGGCTTTGTTGGGCACTCATTG. The threshold cycles for Piezo1/2 transcripts were normalized to three reference genes using the following primers: Tyrosine 3-monooxygenase/tryptophan 5-monooxygenase activation protein zeta polypeptide (YWHAZ): forward CCGTTACTTGGCTGAGGTTG and reverse TGCTTGTTGTGACTGATCGAC; Tata binding protein (TBP): forward TTGGGTTTTCCAGCTAAGTTCT and reverse CCAGGAAATAACTCTGGCTCA; Glyceraldehyde 3-phosphate dehydrogenase (GAPDH): forward AGCCACATCGCTCAGACAC and reverse GCCCAATACGACCAAATCC. YWHAZ and TBP are reported to be stable reference genes for qPCR analysis of MDA-MB-231 derived transcripts^32^.

### Immunostaining and imaging with confocal and total internal reflection fluorescence (TIRF) microscopy

MDA-MB-231 cells were seeded in Nunc Lab-Tek chambers with coverglass bottoms (Thermo Fisher Scientific) and cultured in DMEM Glutamax, 10% FBS. Cell medium was aspirated and the cells were fixed for 10 mins (37°C) with 3.7% formalin, DMEM Glutamax, 10% FBS. The fixative was aspirated and the cell layer was washed extensively with Tris-buffered saline (TBS), Tween 0.1% (TBST). Cells were incubated overnight (4°C) with the specified primary antibodies diluted in TBST. After extensive washing primary antibodies were detected with secondary Alexa Fluor Plus 488 antibodies. Cells were counterstained with phalloidin and DAPI to visualize F-actin and nuclei, respectively. Confocal microscopy of immunofluorescence staining and counterstaining was performed as above on an LSM700 confocal microscope (Zeiss) using a Plan-Apochromat 63x/1.4 (Zeiss) objective and images were captured with Zen software (Zeiss). TIRF microscopy of the same samples was performed on a Nikon TiE microscope equipped with an iLAS2 TIRF illuminator for multi-angle patterned illumination (Cairn Research, Faversham, UK) and a 60x/1.49-NA Apo-TIRF objective (Nikon). Excitation light was delivered by 488-nm and 561-nm diode-pumped solid-state lasers with built-in acousto-optical modulators (Coherent, Santa Clara, USA). Fluorescence was detected with a back-illuminated EMCCD camera (DU-897, Andor Technology, Belfast, Northern Ireland) controlled by MetaMorph (Molecular Devices, San Jose, USA). Emission wavelengths were selected with filters (527/27 nm and 590 nm long-pass, Chroma Technology, Rockingham, USA) mounted in a filter wheel (Lambda 10-3, Sutter Instruments, Novato, USA).

### ERM phosphorylation and phosphatase inhibition assay

MDA-MB-231 cells were seeded into 24-well plates (Sarstedt, Newton, NC, USA) and cultured overnight in DMEM Glutamax, 10% FBS (Thermo Fisher Scientific). Culture medium was aspirated and cells were washed in OptiMEM before being treated as indicated with or without thrombin (1 U/ml) in combination with or without Yoda1 (20 μM) pre-treatment (15 min), co-treatment (5 min) or posttreatment (15 min). Inhibition of phosphatases was performed using the PP1/PP2A phosphatase family inhibitor Calyculin A (Cyc A; 50 nM) or the calcineurin inhibitor Cyclosporin A (Csp A; 250 nM). Cells were stimulated with thrombin (1 U/ml) and after 5 min Cyc A or Csp A was added, 5 min later cells were treated with or without Yoda1 (20 μM). All untreated control conditions were exposed to equal concentrations of DMSO as those present in the Yoda1-, Cyc A- and Csp A-treated cultures.

### SDS-PAGE and Western blotting

MDA-MB-231 cells were treated as indicated and lysates were prepared using RIPA buffer (Thermo Fisher Scientific, Uppsala, Sweden) supplemented with Complete Protease Inhibitor Cocktail (Roche, Basel, Switzerland) and PhosStop (Roche) or Halt (Thermo Fisher Scientific) phosphatase inhibitor cocktail and diluted in 4X Laemmli buffer (BioRad) containing β-mercaptoethanol. Lysates were denatured and proteins were separated by sodium dodecyl sulfate-polyacrylamide gel electrophoresis (SDS-PAGE) on Mini-Protean 10% tris-glycine gels (BioRad). Precision plus protein Kaleidoscope molecular weight (MW) standards (BioRad) were separated in one lane of every gel. Separated proteins were transferred to Immobilon-FL polyvinylidene difluoride (PVDF) membranes (Millipore, Cork, Ireland), which were blocked with a 2:1 ratio of Odyssey blocking buffer (LI-COR Biosciences) and TBS 0.1% Tween-20 (TBST). Membranes were incubated overnight (4°C) with phospho-ERM (pERM) and a-tubulin antibodies. After extensive washing antibodies were detected with secondary goat antirabbit Alexa Fluor 680 IgGs and imaged using an Odyssey Fc system (LI-COR Bioscience). Band intensities were analyzed using Image Studio software (LI-COR Bioscience), and the relative pERM/a-tubulin signal was calculated in Microsoft Excel.

### Statistical analysis

All data were summarized in Microsoft Excel and exported to GraphPad Prism software (GraphPad) for statistical analyses. Statistical significance was tested using two-tailed Student’s t-test, and one-way ANOVA, with Tukey’s multiple comparisons, as indicated in the figures and legends.

## Results

### MDA-MB-231 breast cancer cells exhibit spontaneous blebbing

A subpopulation of MDA-MB-231 cells exhibited spontaneous blebbing when imaged under standard culture conditions (Fig. 1a). F-actin dynamics were visualized by labelling cells with SiR-actin, while changes in cell morphology were simultaneously monitored with differential interference contrast (DIC) microscopy (Fig. 1b-d, Supplementary Video 1). In line with previous reports of bleb dynamics^5,33^, actin-free blebs expanded quickly (within 30 s), followed by actin recruitment and bleb retraction within approximately 2 min (Fig. 1b-d). Z-stack analysis of SiR-actin distribution indicated that blebs formed over the majority of the cell body, but were less prominent at the apical and basal surfaces (Fig. 1e).

**Fig. 1:**
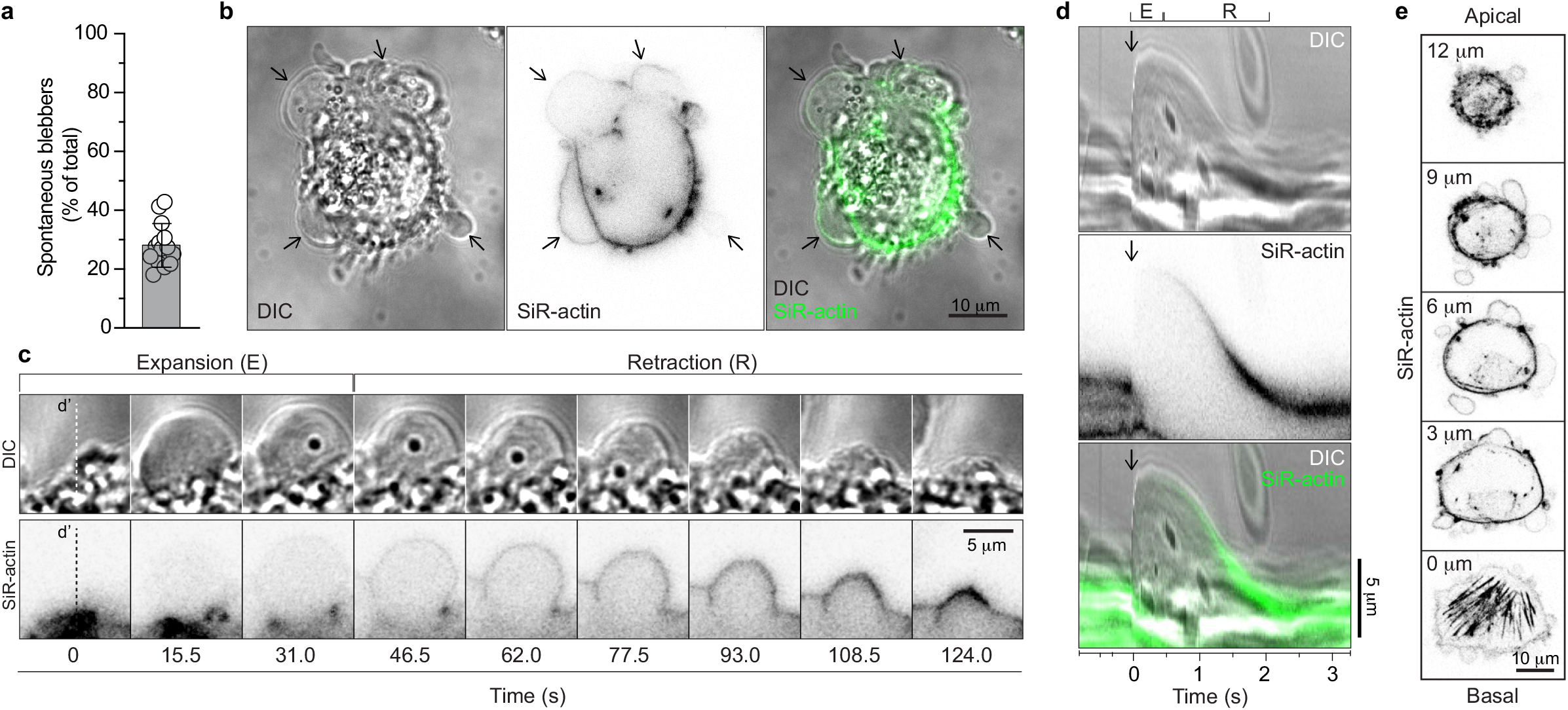
Spontaneous blebbing in MDA-MB-231 cancer cells. **a,** Percentage of spontaneously blebbing MDA-MB-231 cells. Bars represent mean ± s.d. from *n* = 14 separate time-lapse sequences, for a total of 1840 cells. **b,** DIC and confocal microscopy of a blebbing MDA-MB-231 cell with F-actin visualized using the fluorescent SiR-actin probe. Arrows indicate blebs at various stages of F-actin recruitment (Supplementary Video 1). **c,** Time-lapse sequences of the expansion and retraction of a bleb membrane (DIC) and associated F-actin recruitment (SiR-actin) from the cell in (b). **d,** Kymograph of the expansion (E) and retraction (R) of a bleb plotted from the DIC and SiR-actin signals under the dashed lines (d’) in (c). Arrows indicate initiation of bleb expansion. **e,** Confocal z-stack images of the SiR-actin signal from the cell in (b). Positions (μm) in the z-stack are stated relative to the basal side of the cell.

### Thrombin induces blebbing via PAR2 in MDA-MB-231 cells

Tumor environments are typically thrombotic (reviewed in^11^) and thrombin activation of PARs triggers Ca^2+^ release from the ER and this will increase the potential for actomyosin contraction, which is a driving force for bleb formation^21^. Thrombin exposure induced a transient increase in cytosolic Ca^2+^ in MDA-MB-231 cell populations, with similar kinetics observed in individual cells (Fig. 2a-c), and concomitantly increased the percentage of blebbing MDA-MB-231 cells (Fig. 2d). DIC images of thrombin-induced blebbing in a cell is presented in Fig. 2e, f (Supplementary Video 2). Thrombin has the capacity to directly and indirectly activate PAR2^14–16^ and PAR2 expression is elevated in breast tumor specimens and in breast cancer cell lines, including MDA-MB-231 cells^34^. Exposure of MDA-MB-231 cells to the PAR2 agonist SLIGRL mimicked the effects of thrombin with respect to bleb induction and increased cytosolic Ca^2+^ (Fig. 2g-i), with the exception that analysis of individual cell traces revealed that SLIGRL induced Ca^2+^ oscillations (Fig. 2h) that were not typically observed in thrombin-treated cells. DIC images and kymograph of SLIGRL-induced blebbing in a cell are presented in Fig. 2j, k (Supplementary Video 3). Taken together, these results suggest that thrombin-mediated activation of PAR2 is sufficient to induce blebbing in MDA-MB-231 breast cancer cells.

**Fig. 2:**
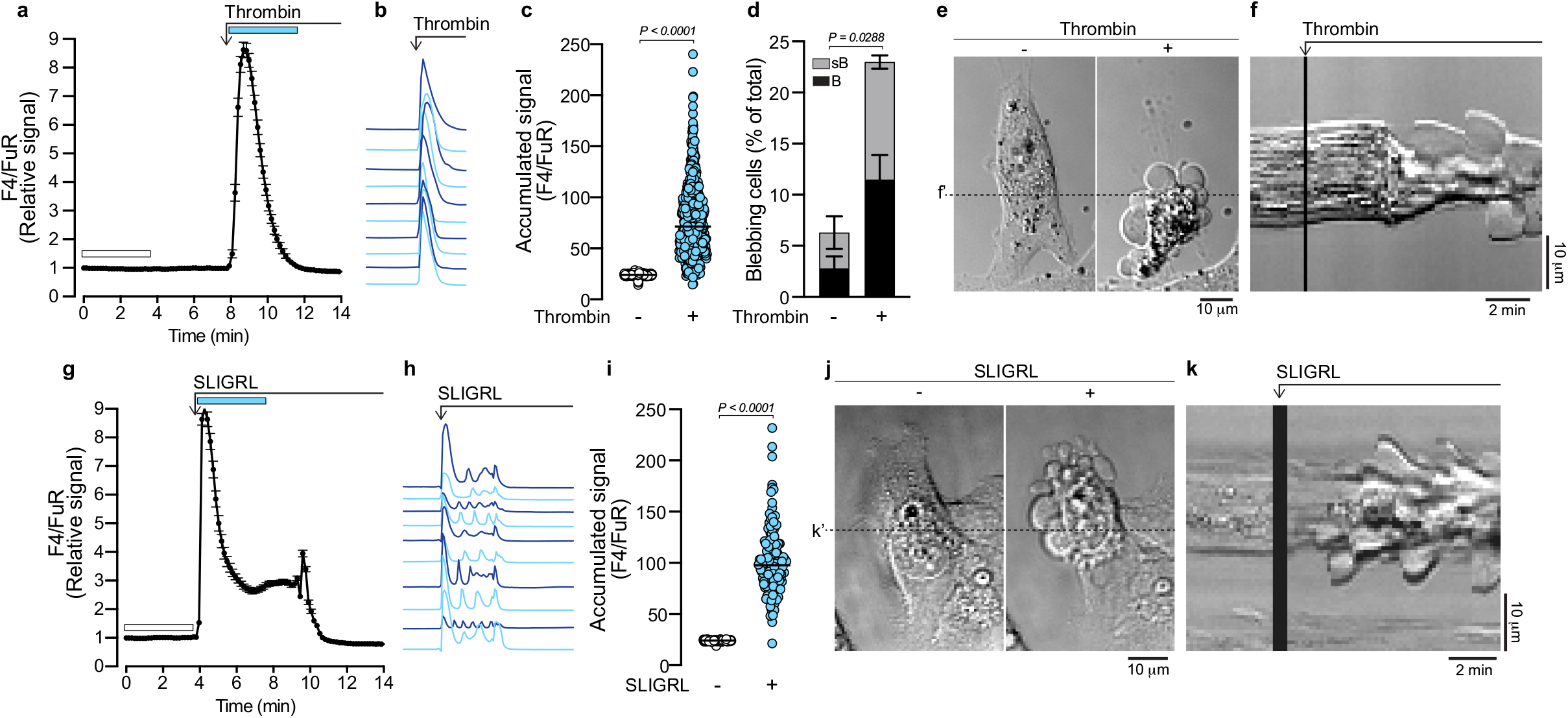
PAR2 activation is sufficient to replicate thrombin-induced blebbing. **a,** Thrombin (1 U/ml) induced changes in cytosolic Ca^2+^ in MDA-MB-231 cells represented by the Fluo-4/Fura Red (F4/FuR) fluorescence ratio recorded by confocal microscopy. The plot represents the F4/FuR signal (mean ± s.e.m.) over time for *n* = 119 cells, from one representative experiment. **b,** Individual F4/FuR signals from ten MDA-MB-231 cells in response to thrombin (1 U/ml). **c,** Accumulated F4/FuR signal in MDA-MB-231 cells before and after thrombin treatment. Scatterplots represent the accumulated F4/FuR signal for the durations indicated by the corresponding color-coded bars in (a) for *n* = 501 cells from three experiments. The */*’-value was determined using the two-tailed Student’s *t*-test. **d,** Quantification of the blebbing status of MDA-MB-231 cells before and after thrombin treatment, determined by analysis of the DIC time-lapse images recorded during the experiments in (c). Bars represent the mean ± s.e.m. B = blebs, sB = small blebs. The *P*-value was determined by two-tailed Student’s *t*-test for percentages of all blebbing cells before and after thrombin addition. **e,** DIC images of an MDA-MB-231 cell before and after thrombin stimulation (Supplementary Video 2). **f,** Kymograph plotted from the DIC time-lapse signal recorded under the dashed line (f’) in (e). **g,** PAR2 agonist (SLIGRL; 10 μM) induced changes in cytosolic Ca^2+^ (F4/FuR ratio) in MDA-MB-231 cells. The plot represents the mean ± s.e.m. over time for *n* = 133 cells, from one representative experiment. **h,** Individual F4/FuR signals from ten MDA-MB-231 cells in response to SLIGRL (10 μM). **i,** Accumulated F4/FuR signal in MDA-MB-231 cells before and after thrombin addition. Scatterplots represent the accumulated F4/FuR signal for the durations indicated by the corresponding color-coded bars in (g) for *n* = 133 cells from one experiment. The *P*-value was determined using the two-tailed Student’s *t*-test. **j,** DIC images of an MDA-MB-231 cell before and after SLIGRL stimulation (Supplementary Video 3). **k,** Kymograph plotted from the DIC time-lapse signal recorded under the dashed line (k’) in (j).

### Contact compression attenuates blebbing in MDA-MB-231 cells

Cellular confinement has been implicated as a driving force in the induction of blebbing behavior^1,8–10^. The capacity of thrombin to induce blebbing in MDA-MB-231 cells (Fig. 2) suggested that such behavior may be adopted by cells in thrombotic tumor environments. To investigate the effects of confinement on blebbing, MDA-MB-231 cells were compressed using a custom-built device called the “Cell Press”, comprised of a PDMS pillar positioned along a linear piezoelectric track that was mounted to a confocal microscope stage (Fig. 3a). Unexpectedly, we observed that spontaneous blebbing exhibited by non-compressed cells was attenuated by gentle contact compression (CC) (Fig. 3b). Further, the attenuation of blebbing by CC was associated with an increase in cytosolic Ca^2+^ (Fig. 3c). Notably, CC did not visibly deform the cells and when subjected to a secondary CC event, which also induced an increase in cytosolic Ca^2+^, the cell’s morphology remained unchanged (Fig. 3b and c). CC proved similarly effective at attenuating thrombin-induced blebbing. Following the thrombin-induced Ca^2+^ response and associated blebbing, the pillar was descended until a clear contact-induced increase in cytosolic Ca^2+^ was observed (Fig. 3d), and the accumulated Ca^2+^ response to CC in the presence of thrombin was significantly greater than that observed for thrombin along (Fig. 3e). DIC images of thrombin-induced, and CC-attenuated blebbing in one MDA-MB-231 cell are presented in Fig. 3f and g (Supplementary Video 4). Notably, even when CC was released, blebbing was not reactivated (Fig. 3g, NC). Quantitative assessment of the blebbing status in MDA-MB-231 populations before and after thrombin treatment, and during subsequent CC confirmed that CC significantly attenuated blebbing in the majority of all blebbing cells (Fig. 3h-i). In line with previous reports, blebbing could be exacerbated in MDA-MB-231 cells when they were subjected to increased compression, which visibly deformed and flattened cells. For example, DIC images revealed that blebbing in a thrombin treated cell was attenuated by CC, but could be reactivated by deformation compression (DC) (Fig. 3j, k; Supplementary Video 5). The kymograph of cytosolic Ca^2+^ changes (F4/FuR fluorescence ratio) for the cell in (j) illustrates that as observed for CC, an increase of cytosolic Ca^2+^ was also observed during DC (Fig. 3j).

**Fig. 3:**
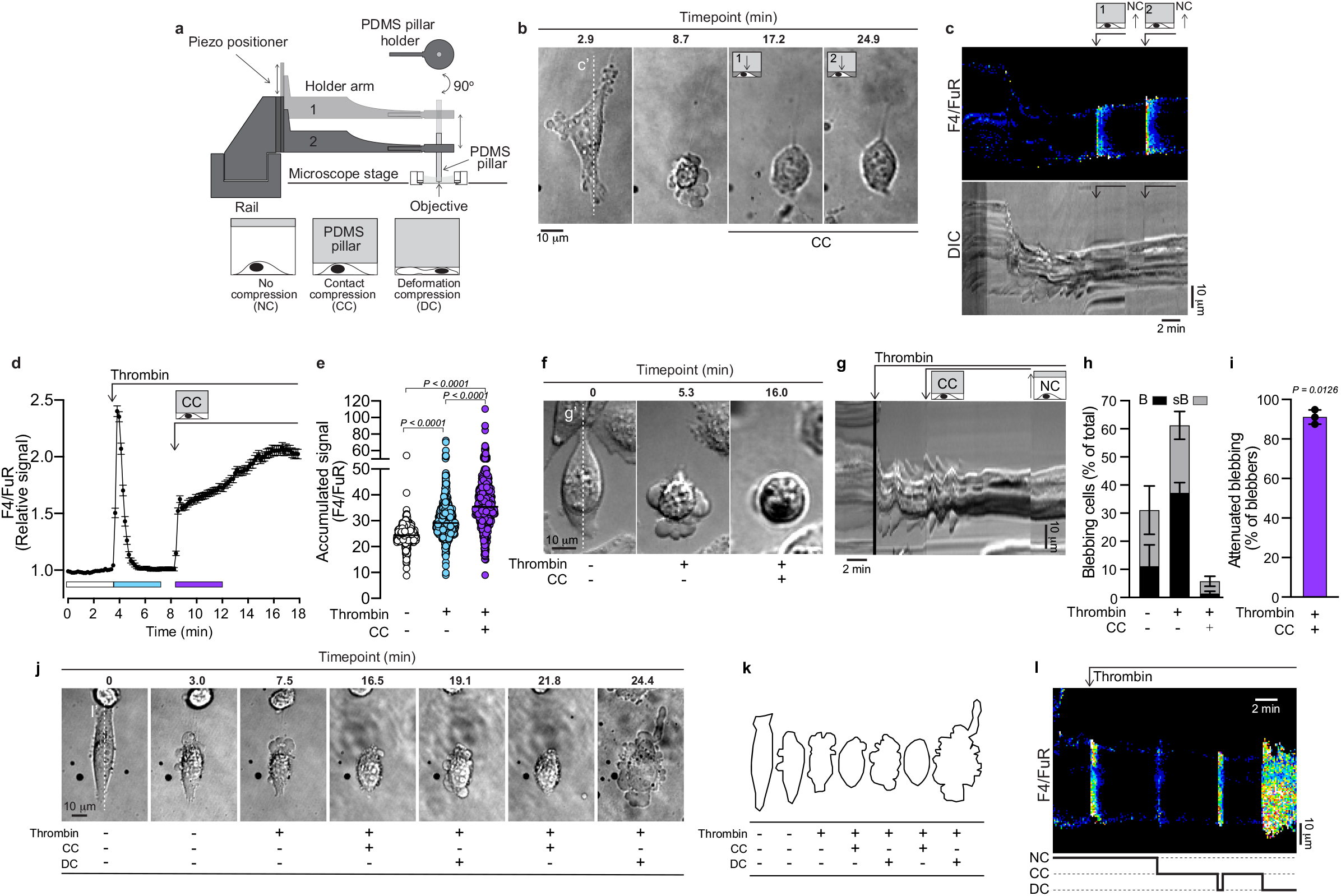
Contact compression attenuates spontaneous and thrombin-induced blebbing. **a,** Overview of the custom-built Cell Press mounted on a confocal microscope stage. MDA-MB-231 cells were subjected to no compression (NC), mild contact compression (CC) or deformation compression (DC) using a PDMS pillar attached to and positioned along a vertical piezoelectric track. **b,** DIC time-lapse images illustrating the induction of spontaneous blebbing and its attenuation by CC in an MDA-MB-231 cell. **c,** Kymographs of the changes in cytosolic Ca^2+^ (F4/FuR ratio) and cell morphology, recorded by confocal and DIC microscopy, respectively. Signals were plotted from under the vertical dashed line (c’) annotated through the cell in (b). The duration of the two CC events are indicated above the images. **d,** Thrombin-(1 U/ml) and CC-induced changes in cytosolic Ca^2+^ (F4/FuR ratio) in MDA-MB-231 cells recorded by confocal microscopy. The plot represents the mean ± s.e.m. over time for *n* = 163 cells from one experiment. **e.** Accumulated F4/FuR signals before and after thrombin, and after CC. Scatterplots represent the accumulated F4/FuR signals for the durations indicated by the color-coded bars in (d), for *n* = 557 cells from three experiments. */*-values were determined by one-way ANOVA, with Tukey’s multiple comparisons. **f,** DIC time-lapse images of an MDA-MB-231 cell illustrating thrombin-induced blebbing and its attenuation by CC. (Supplementary Video 4). **g,** Kymograph of the DIC time-lapse sequence plotted from under the vertical dashed line (g’) in (f). Timepoints for thrombin addition, CC and release of compression leading to NC are annotated. **h,** Quantification of the blebbing status of MDA-MB-231 cells before and after thrombin treatment, and following CC, determined by analysis of the DIC time-lapse images recorded during the experiments in (d). Bars represent the mean ± s.e.m. for the three experiments in (d). (B = blebs, sB = small blebs). **i,** Percentage of cells from (h) in which blebbing was attenuated by CC. Bar represents mean ± s.d. The *P*-value was determined by two-tailed Student’s *t*-test comparing all blebbing cells before and after CC. **j,** DIC time-lapse images of an MDA-MB-231 cell, illustrating the morphological changes exhibited during two cycles of CC followed by DC, in the presence of thrombin (Supplementary Video 5). **k,** Line drawings illustrating the outer perimeter of the cell in (j). **l,** Kymographs of the changes in cytosolic Ca^2+^ (F4/FuR ratio) plotted from under the dashed line (l’) in (j) as recorded by confocal microscopy.

### Characterization of Piezo1 expression and function in MDA-MB-231 cells

The CC-induced Ca^2+^ response, associated with blebbing attenuation (Fig. 3), suggested the activation of a mechanosensitive Ca^2+^ channel, such as Piezo1. Confocal microscopy imaging of cells immunostained for Piezo1 and counterstained to visualize F-actin revealed Piezo1 positive puncta throughout cells (Fig. 4a). To limit our analysis of Piezo1 distribution to the region close to the plasma membrane we employed TIRF microscopy. Piezo1 puncta were again visualized; however, discrete linear aggregates of Piezo1 were also found associated with cortical F-actin filaments (Fig. 4b-d). RT-qPCR revealed Piezo1 mRNA levels to be significantly greater than its related isoform Piezo2 in MDA-MB-231 cells (Fig. 4e). Functional studies of Piezo1 have been facilitated by the chemical agonist Yoda1^35^, which lowers the channels activation threshold^36^. Yoda1 induced a sharp increase in cytosolic Ca^2+^ in MDA-MB-231 cells, which subsequently declined, but typically did not return to baseline for the duration of imaging experiments (Fig. 4f, g). Additionally, the mechanosensitive stretch-activated ion channel inhibitor GsMTx4 effectively impaired the Yoda1-induced Ca^2+^ response (Fig. 4h, i). Together these data confirm the presence of functional Piezo1 in MDA-MB-231 cells.

**Figure 4:**
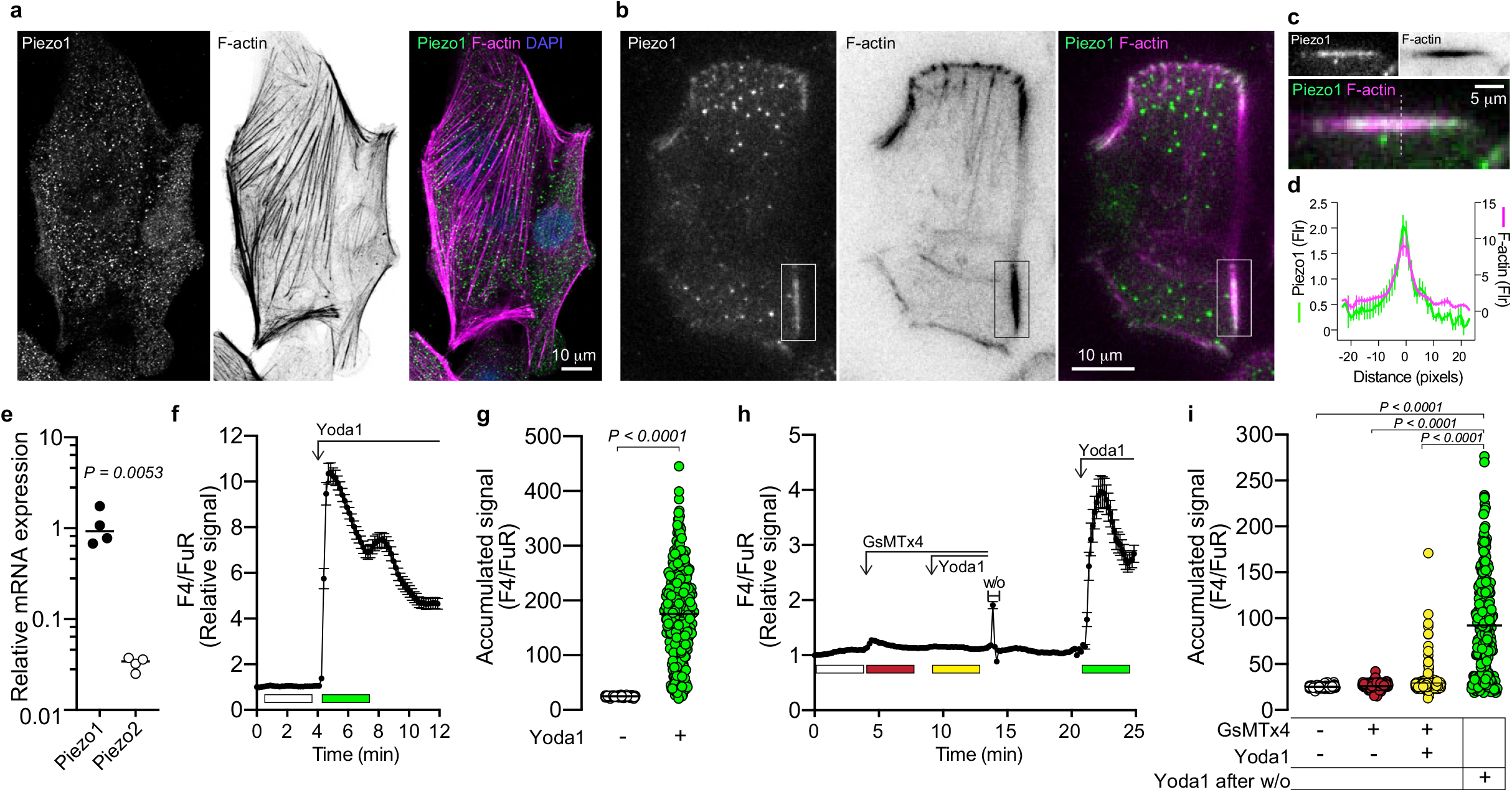
Characterization of Piezo1 in MDA-MB-321 cells. **a,** Confocal microscopy images of Piezo1 immunostaining and counterstaining with phalloidin for Factin, and DAPI for nuclei in MDA-MB-231 cells. **b,** TIRF microscopy images of Piezo1 immunostaining and counterstaining with phalloidin for F-actin in MDA-MB-231 cells. **c,** Enlarged view of the area enclosed by the rectangle in (b). **d,** Relative fluorescence intensity profile of Piezo1 immunosignal and F-actin phalloidin counterstain under linear regions of interest similar to that indicated by the dashed line in (c) i.e. plotted transversely through F-actin filaments. The plot represents the mean ± s.e.m. from 11 structures from three cells. **e,** Relative expression of Piezo1 and Piezo2 mRNA in MDA-MB-231 cells. Scatterplots represent *n* = 4 samples. The *P*-value was determined by two-tailed unpaired Student’s *t*-test. **f,** Yoda1 (20 μM) induced changes to cytosolic Ca^2+^ (F4/FuR signal) in MDA-MB-231 cells recorded by confocal microscopy. The plot represents the mean ± s.e.m. over time for *n* = 148 cells from one experiment. **g,** Accumulated F4/FuR signals before and after exposure to Yoda1 (20 μM). Scatterplots represent the accumulated signals for the durations indicated by the color-coded bars in (f), for *n* = 377 cells from three experiments. The *P*-value was determined by the two-tailed Student’s *t*-test. **h**, Changes to cytosolic Ca^2+^ (F4/FuR signal) in response to Yoda1 (20 μM) in the presence and absence of GsMTx4 (10 μM) recorded by confocal microscopy. The plot represents the mean ± s.e.m. from *n* = 108 cells from one experiment. **i,** Accumulated F4/FuR signals for the durations indicated by the color-coded bars in (h), for *n* = 285 cells from two experiments. The *P*-values were determined by one-way ANOVA, with Tukey’s multiple comparisons.

### The Piezo1 agonist Yoda1 attenuates blebbing

We next assessed the capacity of Yoda1 to attenuate blebbing by activating Piezo1, following thrombin stimulation. To ensure that neither DMSO (in which Yoda1 is dissolved) nor the mechanical stimulation derived from pipetting reagents into the imaging solution were contributing to changes in cytosolic Ca^2+^, a DMSO control solution was added to thrombin-stimulated cells prior to Yoda1 treatment, which had no effect on cytosolic Ca^2+^ (Fig. 5a and b). In contrast, exposure to Yoda1 induced a sharp and sustained elevation of cytosolic Ca^2+^, with an accumulated Ca^2+^ signal that was significantly elevated relative to that induced by thrombin (Fig. 5b). A kymograph of the F4/FuR ratiometric signal and the DIC imaging for a single cell details how the thrombin induced Ca^2+^ increase coincided with the initiation of blebbing, which persisted through the DMSO control, but was markedly attenuated by the addition of Yoda1, which induced a sustained Ca^2+^ increase (Fig. 5c, Supplementary Video 6). Quantitative assessment of the blebbing status in MDA-MB-231 cells confirmed that thrombin increased the percentage of blebbing cells, and this increase was maintained throughout the DMSO control treatment (Fig. 5d). Similar to the effect of contact compression (CC), exposure to Yoda1 attenuated blebbing in the majority of blebbing cells (Fig. 5d and e).

**Figure 5:**
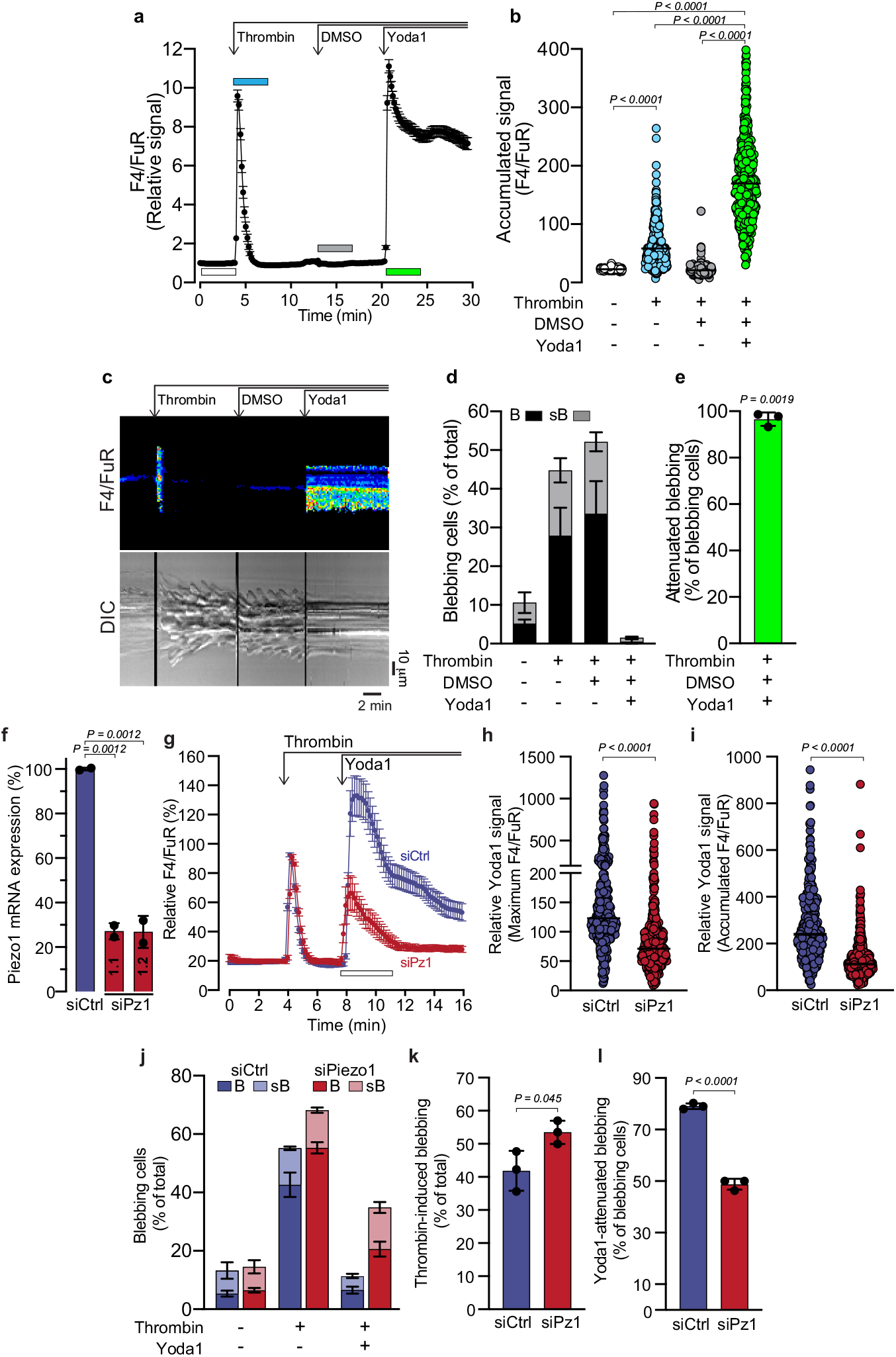
Piezo1 depletion impairs the capacity of Yoda1 to inhibit blebbing. **a,** Thrombin (1 U/ml), DMSO and Yoda1 (20 μM) induced changes to cytosolic Ca^2+^ (F4/FuR ratio) in MDA-MB-231 cells as recorded by confocal microscopy. The plot represents the mean ± s.e.m. over time for *n* = 148 cells from one experiment. **b,** Accumulated F4/FuR signals before and after thrombin, DMSO and Yoda1 addition. Scatterplots represent the accumulated F4/FuR signals for the durations indicated by the corresponding color-coded bars in (a), for *n* = 401 cells from three experiments. *P*-values were determined by one-way ANOVA, with Tukey’s multiple comparisons. **c,** Kymographs of the changes in cytosolic Ca^2+^ (F4/FuR ratio) and cell morphology before and after sequential addition of thrombin, DMSO and Yoda1 of a single MDA-MB-231 cell, recorded by confocal and DIC microscopy (Supplementary Video 6), respectively. **d,** Quantification of the blebbing status of MDA-MB-231 cells before and after sequential addition of thrombin, DMSO, and Yoda1, as determined by analysis of the DIC time-lapse images recorded during the experiments in (b). Bars represent the mean ± s.e.m. B = blebs, sB = small blebs. **e,** Percentage of cells in which blebbing ceased in response to Yoda1; the mean ± s.d. is shown. The *P*-value was determined by two-tailed Student’s *t*-test for percentages of all cells blebbing before and after Yoda1 addition. **f,** Relative expression of Piezo1 mRNA in MDA-MB-231 cells transfected with control siRNA (siCtrl; 50 pM) or siRNA targeting Piezo1 (siPz1: 1.1 and 1.2; 50 pM). Bars represent the mean ± s.d. from two transfections. *P*-values were determined by one-way ANOVA, with Tukey’s multiple comparisons. **g,** Thrombin (1 U/ml) and Yoda1 (20 μM) induced changes to cytosolic Ca^2+^ (F4/FuR ratio) in siCtrl and siPiezo1 transfected MDA-MB-231 recorded by confocal microscopy. The plot represents the mean ± s.e.m. for *n* = 130 siCtrl and *n* = 95 siPz1 transfected cells. **h,** Maximum Yoda1-induced F4/FuR signal relative to maximum thrombin-induced F4/FuR signal in siCtrl and siPiezo1 transfected cells. **i,** Accumulated Yoda1-induced F4/FuR signal relative to accumulated thrombin-induced F4/FuR signal in siCtrl- and siPiezo1-transfected cells. Scatterplots in (h) and (i) represent the maximum and accumulated F4/FuR signals recorded for the durations indicated by the white bar in (g), from *n* = 365 siCtrl cells and *n* = 370 siPz1 cells from three experiments. *P*-values were determined by the two-tailed Student’s *t*-test. **j,** Quantification of blebbing in siCtrl- and siPz1-transfected cells before and after treatment with thrombin, and Yoda1 determined by analysis of the DIC time-lapse images recorded during the experiments in (g); bars represent the mean ± s.e.m. **k,** Percentage of thrombin-induced blebbing in siCtrl- and siPz1-transfected cells. **l,** Percentage of siCtrl and siPz1 cells that stopped blebbing in response to Yoda1. In (j) and (k), bars represent the mean ± s.d. for all blebbing cells from the analysis in (j). *P*-values were determined by the two-tailed Student’s *t*-test.

To determine the effects of Piezo1 loss of function on the Yoda1-mediated attenuation of blebbing MDA-MB-231 cells were transfected with two siRNAs against Piezo1 and RT-qPCR confirmed that Piezo1 transcript expression was significantly reduced (Fig. 5f). Next, MDA-MB-231 cells were treated with siCtrl or both Piezo1 siRNAs (siPiezo1) and then exposed to thrombin followed by treatment with Yoda1. Thrombin- and Yoda1-induced changes in cytosolic Ca^2+^ were plotted for siCtrl- and siPiezo1-treated cells, and for comparison the maximum Ca^2+^ response to thrombin in each experiment was arbitrarily assigned as 100% (Fig. 5g). The maximum and accumulated Ca^2+^ response to Yoda1 was significantly reduced in siPiezo1-treated cells relative to siCtrl conditions (Fig. 5h and i). Quantitative assessment of blebbing status revealed that thrombin-induced blebbing in a slightly greater percentage of siPiezo1-treated cells as compared to siCtrl treated cells (Fig. 5j, k). Importantly, Yoda1’s capacity to attenuate blebbing was significantly reduced in siPiezo1-treated cells as compared to siCtrl-treated cells (Fig. 5l, Supplementary Videos 7, 8). Together these findings suggest that reduced Piezo1 expression renders cells more susceptible to thrombin-induced blebbing, and that Piezo1 activation has the capacity to inhibit or reverse processes that are essential for the maintenance of dynamic blebbing.

### Piezo1 activation attenuates blebbing by suppressing thrombin-induced ERM phosphorylation

To test the possibility that pre-emptive Piezo1 activation could desensitize cells to thrombin induced blebbing MDA-MB-231 cells were pretreated with Yoda1 and then exposed to thrombin. Cytosolic Ca^2+^ was elevated as before in response to Yoda1, while the addition of thrombin was associated with an initial decline in cytosolic Ca^2+^ followed by a gradual increase, before declining again (Fig. 6a). Nonetheless, as demonstrated above, the accumulated Ca^2+^ response elicited by Yoda1 was significantly greater than that for thrombin (Fig. 6b). Quantitative assessment of blebbing status revealed that thrombin failed to increase the percentage of blebbing cells in populations pretreated with Yoda1 (Fig. 6c).

**Figure 6:**
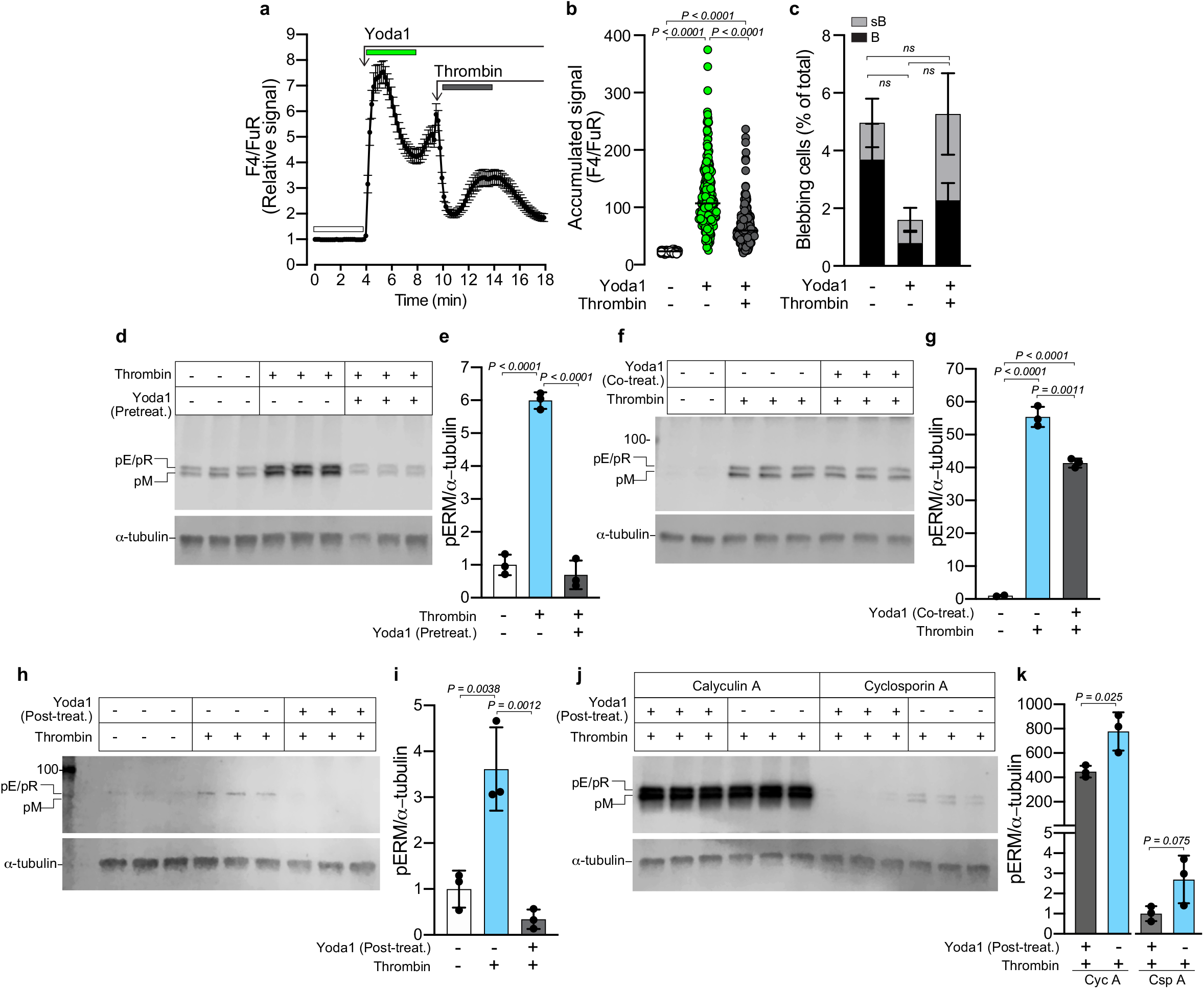
Yoda1 attenuates blebbing by reducing thrombin-induced ERM phosphorylation. **a,** Yoda1 (20 μM) and thrombin (1 U/ml) induced changes to cytosolic Ca^2+^ (F4/FuR ratio) in MDA-MB-231 cells as recorded by confocal microscopy. The plot represents the mean ± s.e.m. over time for *n* = 70 cells from one experiment. **b.** Accumulated F4/FuR signals before and after Yoda1, and subsequent thrombin addition. Scatterplots represent the accumulated F4/FuR signals for the durations indicated by the color-coded bars in (a), for *n* = 258 cells from three experiments. *P*-values were determined by one-way ANOVA, with Tukey’s multiple comparisons. **c,** Quantification of the blebbing status (B = blebs; sB = small blebs) in cells before and after Yoda1, and subsequent thrombin exposure determined by analysis of the DIC time-lapse images recorded during the experiments in (b); bars represent the mean ± s.e.m. *P*-values were determined by one-way ANOVA, with Tukey’s multiple comparisons. **d,** Immunoblotting for phosphorylated ERMs (pERM) in MDA-MB-231 cells pretreated (20 min) with or without Yoda1 (20 μM), followed by thrombin (1 U/ml) exposure (5 min). **e,** Quantification of pERM band intensities relative to a-tubulin loading controls for the samples in (d). Bars represent the mean ± s.d. for three experiments. *P*-values were determined by one-way ANOVA, with Tukey’s multiple comparisons. **f,** Immunoblotting for phosphorylated ERMs in MDA-MB-231 cells treated for 5 min with or without thrombin (1 U/ml) and co-treated with or without Yoda1 (20 μM. **g,** Quantification of pERM band intensities relative to a-tubulin loading controls for the samples analyzed in (f). Bars represent the mean ± s.d. for three experiments, except for the untreated controls where *n* = 2. *P*-values were determined by one-way ANOVA, with Tukey’s multiple comparisons. **h,** Immunoblotting for phosphorylated ERMs in MDA-MB-231 cells stimulated with or without thrombin (1 U/ml), and subsequently (5 min later) treated with or without Yoda1 (20 μM) for 15 mins. **i,** Quantification of pERM band intensities relative to a-tubulin loading controls for the samples in (h). Bars represent the mean ± s.d. for three experiments. *P*-values were determined by one-way ANOVA, with Tukey’s multiple comparisons. **j,** Immunoblotting for phosphorylated ERMs in MDA-MB-231 cells stimulated with thrombin (1 U/ml), followed 5 min later by the addition of Calyculin A (50 nM) or Cyclosporin A (250 nM), and subsequently (5 min later) treated with or without Yoda1 (20 μM) for 15 mins. **k,** Quantification of pERM band intensities relative to a-tubulin loading controls for the samples in (j). Bars represent the mean ± s.d. for three experiments. *P*-values were determined by twotailed Student’s *t*-test.

Phosphorylated ERM proteins tether the actin cytoskeleton to the PM and have previously been shown to be important components of the contractile machinery during bleb retraction^6,7^. Thrombin stimulation (5 min) significantly increased ERM phosphorylation in MDA-MB-231 cells, which was suppressed by pretreating cells (15 min) with Yoda1 (Fig. 6d). Co-treatment of thrombin stimulated cells with Yoda1 (5 min) had a less pronounced effect, but also reduced ERM phosphorylation (Fig. 6 f and g). Further, ERM phosphorylation was similarly reduced in thrombin stimulated cells (5 min) that were subsequently treated (15 min) with Yoda1 (Fig. 6h and i). These data support a role for Piezo1 activation in suppressing or reversing ERM phosphorylation, and provide mechanistic insight into how Piezo1 activation can attenuate blebbing.

A number of phosphatases have the capacity to dephosphorylate ERM proteins including the PP1/PP2A family of serine/threonine phosphatases^37^, which can be inhibited with Calyculin A (Cyc A)^37^. Cells were stimulated with thrombin and after 5 min Cyc A was added, followed 5 min later by treatment (for a period of 15 min) with or without Yoda1. ERM phosphorylation was substantially increased in Cyc A treated cells (Fig. 6j, k), and while band intensity analysis revealed relatively lower ERM phosphorylation in Yoda1 treated samples the levels still far exceeded those observed in experiments with thrombin stimulation alone (Fig. 6d-h). In contrast, addition of cyclosporin A (CspA) to inhibit the phosphatase calcineurin did not produce a similarly robust increase in ERM phosphorylation, and ERM phosphorylation was again lowered in cells treated with Yoda1 (Fig. 6j, k). Together, this implicates the PP1A/PP2A phosphatase family as effectors of Yoda1-stimulated ERM dephosphorylation. Notably, Cyc A alone had no effect on cytosolic Ca^2+^ in MDA-MB-231 cell populations, but did induce blebbing in the majority of treated cells as evident in DIC images from before and after Cyc A treatment (Fig. 7a), suggesting that inhibition of ERM dephosphorylation alone is sufficient to trigger blebbing. As previously, thrombin transiently increased cytosolic Ca^2+^, and cells co-treated with thrombin and Cyc A adopted a blebbing phenotype (Fig. 7b). Finally, and as demonstrated above, Yoda1 attenuated thrombin-induced blebbing in association with elevated cytosolic Ca^2+^ (Fig. 7c); however, co-treatment with Cyc A abolished the bleb-attenuation effects of Yoda1, without effecting the associated elevation of cytosolic Ca^2+^ (Fig. 7d). Together these findings suggest that Piezo1 opening via Yoda1 can activate Ca^2+^-dependent PP1/PP2A phosphatases, which dephosphorylate ERM proteins to attenuate blebbing (Fig. 7e).

**Figure 7:**
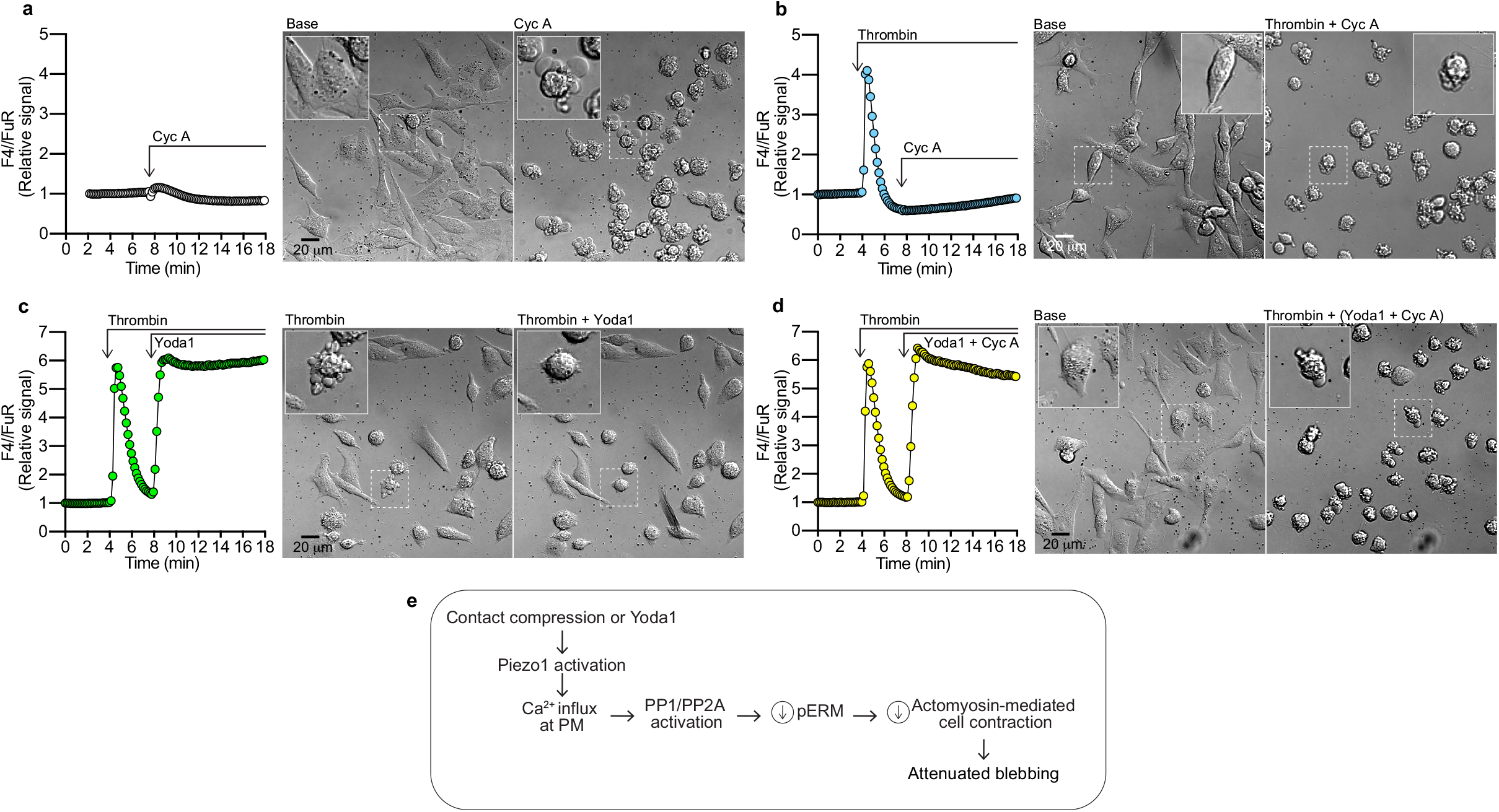
PP1/PP2A phosphatase inhibition induces blebbing. **a-d,** Wholefield analysis of cytosolic Ca^2+^ changes (F4/FuR signal) as recorded by confocal microscopy before and after the indicated treatments with either Calyculin A alone (Cyc A; 50 nM) (a), thrombin (1 U/ml), followed by Cyc A (50 nM) (b), thrombin (1 U/ml) followed by Yoda1 (20 μM) (c) or thrombin (1 U/ml), followed by Yoda1 (20 μM) together with Cyc A (50 nM) (d). Individual frames from DIC time-lapse sequences illustrate cell morphology before and after treatment; insets represent high magnification examples of cells framed with dashed lines in the main images. **e,** Proposed mechanism of Piezo1-mediated attenuation of blebbing. Contact compression or Yoda1-mediated activation of Piezo1 results in an influx of Ca^2+^ at the plasma membrane (PM), activating a Ca^2+^-dependent PP1/PP2A phosphatase, which dephosphorylates ERM proteins, thus reducing the actomyosin-mediated contractile force exerted on the PM, resulting in an attenuation of blebbing.

## Discussion

The plasticity of tumor cells permits them to adopt a range of migratory modes, of which the amoeboid phenotype is defined by plasma membrane blebbing, and is implicated in tumor metastasis^1^. The tumor environment is thrombotic (reviewed in^13^), in part due to the low integrity of tumor vessels, and thrombin has been proposed to increase the invasiveness of MDA-MB-231 breast cancer cells^38^. Here we identify thrombin as a potent stimulus for blebbing in breast cancer cells, and establish that activation of its receptor PAR2 is sufficient to induce this effect. Using the Cell Press device that facilitates controlled compression of cells, we unexpectedly discovered that gentle mechanostimulation through contact compression attenuated blebbing. This bleb attenuation effect was reproduced using the Piezo1 agonist Yoda1, and was compromised in Piezo1-depleted cells. We uncover a mechanism for Piezo1-mediated bleb attenuation, whereby thrombin-mediated ERM phosphorylation is reduced by Piezo1 activation, which is dependent on the activity of PP1/PP2A phosphatases.

The initial expansion of a bleb requires dissociation of the PM from the underlying actin cortex. This actin-free bleb PM expands, after which the contractile machinery required for bleb retraction is sequentially recruited; firstly, the actin-PM cross-linking ERMs, followed by actin and finally myosin II^6^. Increased actomyosin contraction is sufficient to trigger bleb expansion, as it increases hydrostatic pressure within the cell and destabilizes areas of weak actin-PM contact^21^. Consequently, cells deficient in these actin-PM crosslinkers are more likely to bleb, and mechanistic studies of blebbing have been facilitated by the M2 melanoma cell line, which spontaneously bleb in part due to a deficiency in the actin-PM crosslinker filamin^5,39^. Comparatively, the spontaneous blebbing we observe in MDA-MB-231 cells (Fig. 1) is likely explained by their deficiency in the actin-PM linker and tumor suppressor Merlin^40,41^, which is related to the ERM proteins.

Thrombin activation of PARs leads to phospholipase C (PLC)-mediated hydrolysis of PIP2 to DAG and IP3, which likely increases blebbing through a number of pathways. PIP2 serves as a binding site for ERMs; therefore, loss of PIP2 reduces the stability of actin-PM contacts^42,43^, which may promote bleb initiation. IP3-stimulated Ca^2+^ release from the ER will increase actomyosin contraction and drive expansion of actin-free bleb membranes. However, DAG in combination with increased cytosolic Ca^2+^ will also activate protein kinase C (PKC), which can phosphorylate and activate ERMs^44,45^. ERM phosphorylation at the PM exposes their actin-binding domain^46^, and this activation is implicated in coupling the bleb membrane to the actomyosin contractile force that ensures the retraction of actin replenished blebs^6,7^. In the present study, agonist activation of PAR2 was sufficient to reproduce the effects observed for thrombin (Fig. 2). Thrombin has been reported to act directly on PAR2, but may also transactivate PAR2 via the PAR1 tethered ligand^14–16^. Notably, we observed Ca^2+^ oscillations following agonist-activation of PAR2, that were not observed in thrombin treated cells (Fig. 2). It has been proposed that after an initial burst of phosphoinositide hydrolysis, thrombin activation of PAR-tethered ligands is deactivated, while agonist activation of PARs e.g. via SLIGRL does not induce this shutdown^47,48^. This thrombin-specific response would limit the extent of IP3-mediated Ca^2+^ release from the ER, and potentially explains the observed differences in Ca^2+^ dynamics induced by thrombin and SLIGRL. Nonetheless, our results demonstrate that thrombin activation of PAR2 is sufficient to induce blebbing in MDA-MB-231 cells.

Cellular confinement or compression has been implicated as a driving force behind cells adopting the amoeboid mode of migration^4,8,9^. Piezo1 has previously been implicated as a confinement sensor, which suppresses PKA activity and optimizes cell motility in response to the mechanical properties of the environment^27^. Similarly, changes in actomyosin contractility and substrate adhesion can induce cells to transition between bleb- and lamellipodia-dependent cell locomotion modes^2^. Recently, Srivastava et al. identified Piezo in *Dictyostelium* cells as a pressure sensor, required for a Ca^2+^-dependent switch from pseudopod to bleb-based migration^49^. We similarly observed that compressing cells beyond the point of initial contact with the pillar resulted in elevated cytosolic Ca^2+^ and induced blebbing. In contrast, gentle CC attenuated both spontaneous and thrombin-induced blebbing (Fig. 3), as did Piezo1 activation with Yoda1 (Fig. 5). This suggests that the consequences of Piezo1 activation as a result of compression that exceeds some intracellular pressure threshold may be distinct from those associated with Piezo1 activation via Yoda1 or through gentle CC.

The activation threshold of Piezo1 has previously been shown to be lower in cytoskeleton-free PM blebs due to the loss of mechanoresistance that cortical actin provides against changes in membrane tension^30^. Therefore, in blebbing MDA-MB-231 cells, the maximum Piezo1 sensitivity to agonists and mechanostimuli is likely to coincide with the point of maximum bleb expansion, prior to actin recruitment. Borbiro et al. demonstrate that lipid composition also regulates Piezo1, with reduced activity associated with the depletion of phosphoinositides, particularly PIP2^50^. However, while thrombin stimulation will also deplete PIP2, it has been demonstrated that PIP2 is resynthesized within 130 s of PLC-mediated hydrolysis^51^. In our experiments, Piezo1 stimulation (via contact compression or Yoda1) was typically applied no sooner than 4 mins after thrombin exposure and always after cytosolic Ca^2+^ levels had returned to baseline. Therefore, availability of PIP2 is unlikely to limit Piezo1 activity after thrombin stimulation as studied here. Furthermore, while Piezo1 depletion impaired the ability of Yoda1 to attenuate blebbing, Piezo1-depleted cells were also slightly more susceptible to thrombin-induced blebbing (Fig. 5). This suggests that Piezo1 activity may also contribute to counteracting bleb initiation, and this possibility was supported by the fact that thrombin failed to induce blebbing in cells that were pretreated Yoda1 (Fig. 6). Piezo1 is negatively regulated through interactions with the sarcoplasmic/endoplasmic reticulum Ca^2+^ ATPase 2 (SERCA2), and this ER-PM interaction suppresses the migratory capacity of endothelial cells^52^. Bleb expansion may have a negative impact on the integrity of ER:PM contact sites, such as those dependent on actin^53^, and the possibility that this may release SERCA2’s inhibitory effect on Piezo1 warrants further investigation. Taken together, it is likely that the activation threshold for Piezo1 is lower in blebbing than non-blebbing cells. Therefore, the mechanical properties of certain environments may only exceed the Piezo1 activation threshold in cells that are blebbing. Whether this potential for differential activation and bleb silencing represents a general impedance of motility or forces a switch to an alternative mode of migration remains to be evaluated.

ERMs are essential components of the blebbing machinery and their phosphorylation is important for cross-linking actin to the PM, which couples it to actomyosin contractile forces. Thrombin induces ERM phosphorylation in endothelial cells^45^ and a similar effect was observed here in MDA-MB-231 cells (Fig. 6). Importantly, Yoda1 treatment significantly reduced thrombin-induced ERM phosphorylation, offering a molecular explanation for the capacity of Piezo1 activation to attenuate blebbing. TIRF microscopy revealed Piezo1 aggregates that clustered in association with F-actin filaments at the PM (Fig. 4); therefore, it is conceivable that activation of Piezo1 in such clusters would increase the local Ca^2+^ concentration in regions where actin is associated with PM-bound ERM proteins. PP1/PP2A phosphatases are responsible for the dephosphorylation of ERMs in T-cells^37^, and PP1/PP2A inhibition with Cyc A impaired the Yoda1-mediated reduction in phosphorylated ERM levels, which appeared largely unaffected by inhibition of the phosphatase calcineurin (Fig. 6). PP1/PP2A phosphatases can be activated by PKA^54,55^; however, as mentioned above, confinement-induced Piezo1 activation has been associated with reduced PKA activity^27^. Amongst the PP1/PP2A phosphatases the B”/PR72 regulatory subunit is reported to be regulated by Ca^2+^, with two EF-hand motifs that when Ca^2+^ bound activate PP2A phosphatases^56–58^. Based on our data, we propose that Ca^2+^ influx via Piezo1 can activate a PP1/PP2A phosphatase, which dephosphorylates ERMs and in doing so reduces the actomyosin contractile forces exerted on the PM and thereby attenuating blebbing. Cyc A inhibition of PP1/PP2As alone was sufficient to induce blebbing in MDA-MB-231 cells (Fig. 7), indicating that the phosphorylation state of ERMs in MDA-MB-231 cells is in a delicate state of dynamic equilibrium coordinated by the activity of kinases (such as PKC) and PP1/PP2A phosphatases. Any event that tips the balance towards ERM phosphorylation is therefore likely to increase the likelihood of blebbing. PP2As are important tumor suppressors; however, in breast tumors and in breast cancer cell lines, including MDA-MB-231 cells, endogenous PP2A inhibitors are overexpressed^59^. In the light of our findings, downregulation of PP2As in MDA-MB-231 may also contribute to their susceptibility to blebbing, including thrombin-induced blebbing, while Piezo1-mediated activation of PP2As dephosphorylates ERMs and impedes bleb initiation or suppresses ongoing dynamic blebbing. The ability of thrombin to induce blebbing, and Piezo1 to attenuate blebbing, identifies them as potential targets for impairing the capacity of cancer cells to adopt dynamic blebbing behavior associated with the amoeboid phenotype, and future studies are warranted to determine their impact on the risk of metastasis.

## Contributions

P.O’C. and J.K. conceived of the study and designed the experiments. P.O’C. performed the experiments, acquired and analyzed the data, and visualized the results. P.O’C., G.S., O.I-H., and J.K. discussed data interpretation. A.E. designed, produced and assembled the “Cell Press”, with modifications performed by P.O’C, and N.F.K. G.S. assisted with TIRF microscopy. P.O’C., O.I.H., and J.K. acquired funding. O.I-H., and J.K. provided resources. P.O’C. and J.K. drafted the original manuscript, and A.E., N.F.K., G.S., and O.I-H. contributed edits and revisions. All authors approved the submitted manuscript.

## Acknowledgements

This study was funded by grants to J.K. from Cancerfonden (CAN 2017/703), to P.O’C. from O.E. och Edla Johanssons vetenskapliga stiftelse, and to O.I-H. from the Swedish Research Council (MH2015-03087), and the Göran Gustafsson Foundation. 3D printing was performed at U-PRINT: Uppsala University’s 3D-printing facility at the Disciplinary Domain of Medicine and Pharmacy.

