## Supplementary Video Legends for "Piezo1 activation attenuates thrombin-induced blebbing in breast cancer cells"

*Corresponding authors

**Supplementary Video Legends**

**Supplementary Video 1.**

**Plasma membrane and actin dynamics in a spontaneously blebbing MDA-MB-231 cell.** Time-lapse series of DIC and confocal microscopy recordings for the blebbing MDA-MB-231 cell in Fig. 1a with F-actin visualized using the fluorescent SiR-actin probe. The format of the digital timer is min:sec.msec. Scale bar; 10 μm.

**Supplementary Video 2.**

**Thrombin-induced blebbing in an MDA-MB-231 cell.** DIC time-lapse recordings of thrombin-induced blebbing for the MDA-MB-231 cell in Fig. 2e, f. The addition of thrombin (1 U/ml) is indicated as +Th. The format of the digital timer is min:sec. Scale bar; 10 μm.

**Supplementary Video 3.**

**SLIGRL-induced blebbing in an MDA-MB-231 cell.** DIC time-lapse recordings of SLIGRL-induced blebbing for the MDA-MB-231 cell in Fig. 2j, k. The addition of SLIGRL (10 μM) is indicated as +SLIGRL. The format of the digital timer is min:sec. Scale bar; 10 μm.

**Supplementary Video 4.**

**Contact compression attenuates thrombin-induced blebbing in an MDA-MB-231 cell.** DIC time-lapse recordings of contact compression (CC) attenuation of thrombin-induced blebbing for the MDA-MB-231 cell in Fig. 3f, g. The addition of thrombin (1 U/ml) is indicated as +Th, and the application of contact compression as +CC. Removal of the compression pillar is indicated as NC (no compression). The format of the digital timer is min:sec. Scale bar; 10 μm

**Supplementary Video 5.**

**Deformation compression of an MDA-MB-231 cell is associated with blebbing.** DIC time-lapse recordings of deformation compression (DC) leading to exaggerated blebbing in the MDA-MB-231 cell from Fig. 3j-l. The addition of thrombin (1 U/ml) is indicated as +Th, and the application of contact compression or deformation compression are indicated as +CC and +DC, respectively. The format of the digital timer is min:sec. Scale bar; 10 μm

**Supplementary Video 6.**

**Yoda1 attenuated blebbing in a thrombin-treated MDA-MB-231 cell.** DIC time-lapse recordings of the MDA-MB-231 cell from Fig. 5c. The addition of thrombin (1 U/ml), DMSO and Yoda1 (20 μM) are indicated as +Th, +DMSO, and +Yoda1. The format of the digital timer is min:sec. Scale bar; 10 μm

**Supplementary Video 7.**

**Yoda1 attenuation of thrombin-induced blebbing in an siCtrl-treated MDA-MB-231 cell**

DIC time-lapse recordings of an siCtrl-treated MDA-MB-231 cell. The addition of thrombin (1 U/ml), followed by Yoda1 (20 μM) are indicated as +Th and +Yoda1. The format of the digital timer is min:sec. Scale bar; 10 μm

**Supplementary Video 8.**

**Failure of Yoda1 to attenuate thrombin-induced blebbing in an siPiezo1-treated MDA-MB-231 cell.** DIC time-lapse recordings of an siPz1-treated MDA-MB-231 cell. The addition of thrombin (1 U/ml), followed by Yoda1 (20 μM) are indicated as +Th and +Yoda1. The format of the digital timer is min:sec. Scale bar; 10 μm
